# Tolerance to herbicide drift in “weedy” species is context-dependent and driven by early vegetative growth

**DOI:** 10.64898/2026.09.14.751410

**Authors:** Veronica Iriart, Tia-Lynn Ashman, Matthew R. Armstrong, Sara Colom, Toshiro Newsum, Anah Soble Botran, Regina S. Baucom

## Abstract

While plant communities that exist at the edges of agricultural fields are typically composed of “weedy” species, they provide essential resources for insect pollinators and other wildlife. However, little is known about how sublethal exposures to herbicides via drift (off-target chemical movement by air) affects the fitness of weedy plant species across varying agro-ecological environments. If weed species vary in sensitivity to herbicide drift, then this could result in plant community shifts in favor of more tolerant species. Consequently, these changes in species composition could alter resource availability in agroecosystems. Here, we replicated a common garden experiment in three US states where agriculture is a major form of land use (Michigan, Pennsylvania, and Tennessee). We exposed nine weed species to a sublethal, drift-level rate of the herbicide dicamba and examined plant fitness. We found that fitness in the presence of dicamba drift depended on both the weed species and the environmental context. For some species, dicamba drift significantly reduced fitness. For others, it had no effect, indicating greater tolerance to the herbicide. However, more variation in tolerance to dicamba drift among species was observed in Michigan compared to Pennsylvania or Tennessee, a result that was partially explained by the severity of damage from insect herbivory that occurred at each location. In a follow-up study, we explored the relationships between plant developmental traits and fitness to determine the functional pathways that underlie tolerance to dicamba drift using structural equation modeling. This approach revealed that weedy plants that are better able to buffer the effects of dicamba drift on short-term growth are then better equipped to minimize effects on flower production and ultimately fitness (seed set). Overall, our work advances knowledge about how wild plant populations in different contexts are responding to unintentional, yet increasingly common occurrences of anthropogenic stress in the form of off-target herbicide exposures.

**Open Research Statement:** Data and code used to generate analyses and results are publicly available for review at https://github.com/iriav1418/WeedShiftsProj.

## Introduction

Environmental change caused by human activity can represent a turning point for wild populations. These disruptions can dramatically alter population demographics, community assembly, and species interactions (Johnson & Munshi-South, 2017; Valiente-Banuet et al., 2015). Given that half of the world’s land is currently used for agriculture (FAO, 2024; Poor & Nemecek, 2018: Richie & Roser, 2024), new pesticides and other agrochemicals are especially likely to become novel selective agents that catalyze major community shifts in global ecosystems (e.g. Bohnenblust et al., 2014). In particular, a long-standing issue regarding the use of herbicides for weed control is their likelihood to “drift”, or move through the atmosphere at low concentrations (0.1-5% of the field application rate) and onto non-target areas (Cessna et al., 2005, Bohnenblust et al., 2016). However, from an ecological standpoint, little is known about how occurrences of herbicide drift affect the wild systems that are most vulnerable to this source of anthropogenic stress–i.e., crop-associated (“weedy”) plant communities (Iriart et al., 2020). Despite being broadly regarded as pests, weed communities in unmanaged areas near agricultural land provide important ecosystem services, namely by supporting plant biodiversity and producing essential resources for insect pollinators throughout the growing season (Bretagnolle & Gaba, 2015; Ouvrard et al., 2018). Therefore, illuminating the ways in which herbicide drift exposures alter the composition, ecology, and/or evolution of weed communities remains a pertinent environmental concern, especially as agricultural practices continue expanding and intensifying.

One key mechanism by which herbicide drift could reshape naturally-occurring weed communities is through interspecific variation in tolerance to herbicide stress. For example, weed species that are more herbicide-tolerant would be expected to remain or increase in abundance, while non-tolerant species decline or even go extinct. Similar shifts in plant communities driven by variation in tolerance to other stressors (including other agricultural management practices) have been well-documented (Ghersa et al., 1994; Johnson et al., 2009; Swanton et al., 1993). However, the potential for such community shifts to occur due to interspecific tolerances has yet to be linked to foundational studies in the field of evolutionary ecology that examine tolerance as a fitness outcome. In these studies, tolerance is the ability of an organism to minimize the negative effects of fitness when exposed to a stressor (Baucom & Mauricio, 2004, 2008; Fineblum & Rausher, 1995). A species that shows no difference in fitness when in the presence of a stressor compared to in its absence is considered completely tolerant. Meanwhile, species that have significantly lower fitness in the presence of a stressor are characterized as undercompensating, and species that reach significantly higher fitness under stress are classified as overcompensating (Baucom & Mauricio, 2004). In characterizing weed species’ tolerances to herbicides in this way, we would be better equipped to understand contemporary changes in the composition of these plant communities that interface with agriculture and possibly predict their evolutionary trajectories.

A small set of previous studies have indicated that common weedy plant species indeed show significant variation in tolerance to drift from various herbicides, which could lead to these proposed changes in community composition (Iriart et al., 2022; Baucom et al., 2025; Olszyk et al., 2017). Many of these species have been shown to undercompensate for or tolerate herbicide drift, but a minority have shown overcompensatory responses. However, it is yet unknown whether they show plasticity in tolerance across field environments. Abiotic climatic variables can induce changes in plant physiology that could affect how plants respond to an herbicide (Ziska, 2016), thus leading to plastic responses. For example, higher temperatures can reduce stomata or increase leaf thickness, and this could reduce chemical absorption, making some weedy plants more herbicide-tolerant (Ziska & Bunce, 2006). Further, biotic variables, particularly the intensity of herbivory, may also mediate the effects of herbicides on plant fitness (Johnson et al., 2019), and thereby affect herbicide tolerance. In a prior field experiment (Johnson et al., 2022), it was found that the proportion of chewing damage caused by herbivory increased in *Datura stramonium* and *Abutilon theophrasti* when plants were exposed to a drift-level rate of the herbicide dicamba compared to unexposed controls, but the amount of herbivory on *Ipomoea purpurea* was unaffected. Nevertheless, despite clear knowledge that the agricultural matrix encompasses a wide range of abiotic and biotic conditions (Potapov et al., 2021), no study has previously explored the effects of herbicide drift on weedy plant communities across realistically variable agro-eco environments.

Moreover, the functional pathways that could indicate whether a weed species demonstrates complete tolerance, undercompensation, or overcompensation to herbicide drift remain poorly understood, yet these would be valuable for predicting community shifts. Past work has shown that, for undercompensating weedy plants, low-dose exposures to herbicides may cause vegetative deformities, impede growth, delay flowering, reduce floral displays, and decrease pollinator visitation (Baucom et al., 2025; Bohnenblust et al., 2016; Iriart et al., 2022; Iriart et al., 2024; Johnson et al., 2022; Olszyk et al., 2017). While some of these studies have also characterized the effect of herbicide drift on plant fitness, none have directly explored relationships between these phenotypic responses to herbicide exposure (i.e., changes in vegetative growth or flowering) and fitness. For instance, the ability of a plant to recover vegetative growth shortly after being exposed to herbicide drift might result in a fitness cost, where limited resources are allocated towards short-term rather than long-term development (Baucom & Mauricio, 2004). Alternatively, significant alterations in flowering onset and/or flower production after drift might strongly relate to fitness if these changes result in a lesser or greater number of beneficial plant-pollinator interactions. In uncovering these relationships, we would gain a greater appreciation of how herbicide exposure directly affects plant fitness (e.g. by influencing fitness metrics like final biomass or seed production) and/or indirectly affects plant fitness (e.g. by influencing traits that directly affect fitness metrics). Simultaneously, we would learn more about the potential downstream ecological consequences of these herbicides, especially for pollinators whose populations are bolstered by the presence of certain flowering phenotypes (Cappellari et al., 2022; Fornoff et al., 2016).

In this study, we investigated whether weedy plant species varied in their tolerance to herbicide drift and whether such tolerance depended on the field environment. We also asked which traits—leaf morphology, vegetative growth, flowering phenology, and flower number—contribute to variation in fitness under herbicide exposure, thereby shaping tolerance to herbicide drift. We used a sublethal dose of the synthetic auxin herbicide dicamba that simulated an occurrence of herbicide drift to address these questions. Of the various classes of herbicides used in agriculture today, synthetic auxin herbicides are particularly likely to drift across agricultural landscapes (Mortenson et al., 2012). These herbicides act by mimicking the plant growth hormone auxin, causing aberrant growth in dicot plants (Todd et al., 2020). Both 2,4-D and dicamba are not new herbicides–they were first commercialized in the 1940s and 1960s, respectively (Egan et al., 2014). However, over the last decade, the development of 2,4-D-and dicamba-tolerant crops has resulted in the rapid, wide-spread adoption of these herbicides (especially dicamba), causing many wild communities of weedy plants to be newly exposed to these chemicals (Canella Vieira et al., 2023; USGS, 2024a; USGS, 2024b).

We conducted two complementary summer field experiments to achieve these goals. In the first field experiment performed in 2019, we examined the effects of dicamba drift on plant fitness across nine common weedy species planted in three different environmental gardens located in Michigan, Pennsylvania, and Tennessee, USA. These geographic areas represent similar environments based on past dicamba use (USGS, 2019), but diverge in their summer climates (NOAA, 2026), and potentially levels of herbivory (Meineke et al., 2018). Specifically, we answered the following questions: (1) To what extent do weed species vary in tolerance to dicamba drift exposure? (2a) Does species’ tolerance to dicamba drift vary depending on the environmental context (i.e., geographic location)? (2b) If so, could the intensity of herbivory or climate explain the variation in dicamba tolerance observed? In the second field experiment performed in 2021, we investigated a similar set of weed species in the Michigan garden, re-examined the effects of dicamba drift on fitness (question 1), and explored whether findings of interspecific variation in dicamba drift tolerance were replicated across field experiments conducted in the same location. In this experiment, we additionally recorded various growth and flowering responses to dicamba drift exposure, and used structural equation modeling to elucidate the direct and indirect effects of dicamba drift on plant fitness. We then answered: (3) Which functional traits (e.g. vegetative and/or flowering traits) of weedy plants underlie the effect of dicamba drift on plant fitness and thus dicamba drift tolerance?

## Materials and Methods

### Seed Collection

We sampled seeds from 26 species of naturally-occurring weed communities at the edge of soybean, maize, or fallow fields located in central Tennessee or western Kentucky in 2018 and 2019 and used these seeds in the field experiments performed in 2019 and 2021. Mature seeds were collected from 1-3 populations per species and mixed in bulk to ensure that seed origin was randomized in field experiments. Location and population information of seed sources of all species that were planted can be found in Appendix S1: Table S1 and Appendix S1: Table S2 for the 2019 experiment and in Baucom et al. (2025) for the 2021 experiment. Agricultural use of dicamba in both areas of Tennessee and Kentucky where seeds were collected began in 2017 and was predicted to increase (personal communication with local extension agents, Baucom).

### 2019 Field Experiment

To determine whether tolerance to dicamba drift varied according to plant species and the environmental context, we planted three seeds of 26 common weed species (Appendix S1: Table S1) into tilled soil within 11 (4 × 2 m) plots surrounded by deer fencing at each of the three experimental common gardens. These were located in the Matthaei Botanical Gardens in Ann Arbor, Michigan (42.3024°N, 83.6632°W), the Pymatuning Laboratory of Ecology in Linesville, Pennsylvania (41.6433°N, 80.4280°W), and at a 10 acre private residence in Apison, Tennessee, USA (35.0206° N, 85.0205° W) (hereafter, the “Michigan”, “Pennsylvania”, and “Tennessee” gardens, respectfully). In total, 858 seeds (i.e., 3 x 26 x 11) were planted in 2019 on May 16 in Pennsylvania, May 26 in Tennessee, and June 3 in Michigan. In both the Michigan and Pennsylvania gardens, one species, *Cardiospermum halicacabum*, was germinated in the greenhouse and transplanted into plots. Nine species consistently established six or more individuals across plots in Michigan and Pennsylvania, and thus were considered “focal species” (Table 1). In Tennessee, germination of all but four focal species was very low due to drought and high heat and re-plantings were not attempted given the severity of local conditions.

**Table 1.**
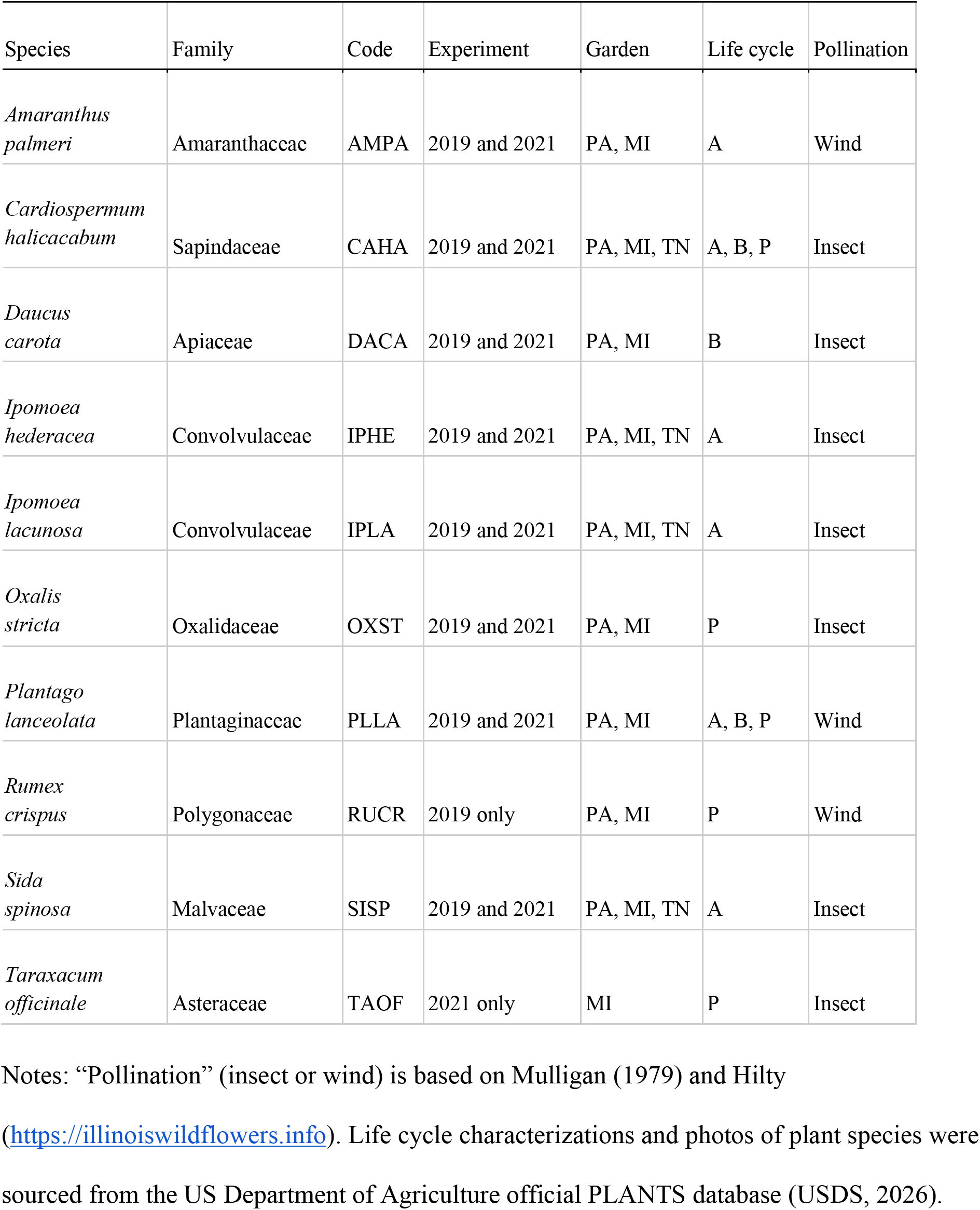
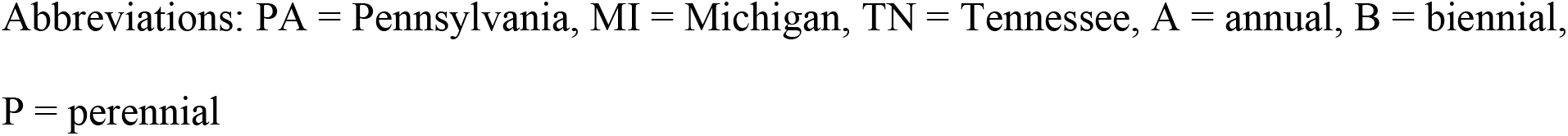
Focal weedy plant species investigated in the 2019 and 2021 field experiments.

Three weeks post-germination, we applied herbicide treatments to experimental plants. We applied 0.7% of the field application rate of dicamba (561 g active ingredient (3,6-dichloro-o-anisic acid) per hectare; Albaugh, LLC, Ankeny, IA; Albaugh, 2018) to plants within five randomly chosen plots, and a 0% control to another five plots. The 0.7% dicamba treatment is within the range of dicamba concentrations that are representative of herbicide drift (Bohnenblust et al., 2016; Cessna et al., 2005). We applied the field application rate to the final plot per garden. We included “Preference” surfactant (non-ionic surfactant blend, WinField Solutions, St. Paul, MN) in all herbicide treatments at 0.1% v/v. To apply treatments, handheld multipurpose sprayers (Model #1501HDX; HD Hudson Manufacturing, Lowell, MI) were used in the Michigan and Tennessee gardens, and a diaphragm pump backpack sprayer (Model #473-D[VI1], Solo Inc., Newport News, VA, USA) was used in the Pennsylvania garden. At all gardens, the sprayers were set to a medium-fine mist with an operating pressure of 40-45 PSI to spray plants with one pass until just wet. Immediately prior to treatment application, we recorded the length of the largest leaf in mm with a digital caliper, counted the total number of leaves, and used the product to estimate pre-treatment plant size of each plant. At three weeks post-treatment, we re-measured plant size and recorded survival to evaluate the efficacy of the dicamba treatments. Overall, 1-14 replicate plants (mean = 8) were treated with either the dicamba drift treatment or control per focal species per garden (total sample size = 464 plants), and 1-7 replicate plants (mean = 4) were treated with the full-dose treatment per focal species (excluding *Daucus carota* that was absent in full-dose plots) across the three gardens (total sample size = 32).

Although deer fencing protected plants from herbivory caused by large herbivores, experimental plants were still vulnerable to herbivory by insects, which can also significantly affect plant biomass (Carson & Root, 1999). Consequently, we investigated whether patterns of insect herbivory could explain variation in plant biomass and dicamba tolerance (question 2b). Specifically, we surveyed plants for insect herbivory at three weeks post-treatment. We categorized each plant into one of three levels of herbivory (zero/minimal, moderate, or severe) based on visual assessments of the amount of missing tissue apparent on the whole plant.

At ∼70 days post-treatment in each garden (September 5 in Pennsylvania, September 15 in Tennessee, and September 25 in Michigan), we harvested the above ground vegetation of all surviving plants. We dried shoots at 70°C for three days, and used this measure of shoot biomass as a rough estimate of plant fitness (Younginger et al., 2017). We acquired climate data from the closest weather station to each garden: the Ann Arbor Municipal Airport in Michigan, the Meadville Port Meadville Airport in Pennsylvania, and the Chattanooga Airport in Tennessee (NOAA, 2026). We used this data to calculate the average daily temperature and precipitation while plants were growing in each garden. For days when precipitation was recorded as “trace amounts” (< 2.54 mm), we added a minimum positive number (0.254 mm) to the dataset.

### 2021 Field Experiment

To better understand the effects of dicamba drift on weedy plant species and identify which plant functional traits underlie tolerance to dicamba drift, we performed a second field experiment. Details of this experiment, including a complete list of the species that were planted, were outlined previously (Baucom et al., 2025). In brief, we planted six replicate seeds of 11 common weed species within 11 plots at the Michigan garden on June 1, 2021. Replicates of each species were likewise planted in the greenhouse at this time for transplanting; since germination in the field was low, we transplanted two to four replicates of each species into each of the field plots on June 22-23. Preliminary data analysis indicated no effect of transplantation and as such this term was not retained in subsequent analyses. Four weeks post-transplantation, five randomly-selected plots were treated with 0.7% dicamba, four others were treated with 0% dicamba, and the final two plots were treated with the field application rate of dicamba using a handheld sprayer as described above. As before, we estimated pre-treatment plant size prior to applying dicamba.

To determine functional traits that may mediate the effects of dicamba exposure on plant fitness, we measured two early growth responses to dicamba drift (plant size and leaf damage) and tracked flowering. Specifically, at three weeks post-dicamba treatment, we recorded the total number of leaves and the length of the largest leaf (mm) and used the product of these to estimate early vegetative plant size (3wk size), and we recorded the number of leaves showing signs of dicamba damage (leaf damage). Typically, this was noted by the presence of leaf “cupping”, where leaves are wrinkled and take on a cupped shape (Johnson et al., 2023). From July 23 to September 29, we recorded the day of first flowering for each individual and the number of flowers present per plant three times a week.

To characterize dicamba drift tolerance in this experiment, we used seed number to assess differences in plant fitness in drift-treated vs. control plants, as seed number is a more typical measure of fitness in tolerance studies (Agrawal et al., 1999; Baucom & Mauricio, 2004; Kover & Schall, 2002; Mauricio et al., 1997). To do so, we sampled fruits that each flowering individual produced over the course of the field experiment and used them to estimate the total number of seeds produced. A couple of flowering *Amaranthus palmeri* plants (N = 2/40) were identified as male and thus were excluded from the data set. We counted the number of seeds produced per a random sample of 50 fruits per species and then performed a regression to estimate total seed count per plant. Nine of the 11 species produced sufficient fruit to measure seed number in this way, so we identified these as the focal species of this experiment (Table 1). Overall, the number of replicate plants ranged from 2-22 (mean = 12) per treatment (control and drift) per focal species for a total sample size of 219 plants.

### Data Analysis

We performed all analyses in R version 4.2.2 (R Core Team, 2026). We used the *lme4* package (Bates et al., 2015) to run mixed-effects and generalized mixed-effects linear models. We modeled data from plants that had received the control and drift treatments only. After inspecting residuals and validating models using the *DHARMa* package (Hartig, 2022), we tested the significance of fixed effects with Type III sums of squares using the *Anova* function from the *car* package (Fox & Weisberg, 2019). We calculated estimated marginal means, tested contrasts, and conducted pairwise comparisons as described below using the *emmeans* package (Lenth, 2025). We used the ggplot2 package to create all figures (Wickham, 2016).

Specifically, using data from the 2019 field experiment, we answered questions 1 and 2a by constructing the “Fitness Model”:

Fitness ∼ treatment × species × garden + pre-treatment plant size + (1|plot)

where fitness was input as shoot biomass data (natural log-transformed). Treatment, species, and garden (Michigan, Pennsylvania, and Tennessee) were included as fixed effects, plot was included as a random effect, and pre-treatment size was included as a covariate. A significant interaction between treatment and species would indicate that species varied in the way that dicamba drift exposure affected their fitness, i.e. species within the weed communities varied in tolerance to dicamba drift, in accordance with previous findings (e.g., Iriart et al., 2022). A significant three-way interaction between treatment, species, and garden would provide new insight to suggest that species’ tolerance to dicamba drift depended on the environmental context in which weed communities were based.

When we found a significant treatment x species interaction, we determined tolerance to dicamba among species by examining species-level biomass in both the presence and absence of dicamba drift. Specifically, we calculated contrast estimates (Abdi & Williams, 2010) for each species, which reflects the difference in the estimated marginal means for biomass between treatments and identified species estimates that were significantly different from zero. As described previously, tolerance is commonly defined as the maintenance of fitness in the presence of the damaging agent (Fineblum & Rausher, 1995). Therefore, we considered species that showed significant fitness reduction in the presence of dicamba drift, relative to fitness in the absence of dicamba drift, as exhibiting low tolerance to dicamba drift. Conversely, we characterized species that showed no reduction in fitness as tolerant to dicamba drift. If we also found support for a significant treatment x species x garden interaction, then we examined patterns regarding species’ tolerance according to the different gardens.

We also examined whether patterns of herbivory varied according to the different experimental conditions (garden, plant species, and herbicide treatment) in the 2019 study. Because only 5% of experimental plants were categorized as having experienced “severe” herbivory, to improve model performance and generalization, we analyzed herbivory as a binary variable. Plants were assigned a 0 if they were recorded as having zero/minimal herbivory or a 1 if they were recorded as having moderate or severe herbivory. We then ran a generalized linear mixed effects model with a binomial distribution using herbivory as the response variable, and the main effects of garden, plant species, and herbicide treatment as explanatory variables. We conducted *post-hoc* pairwise comparisons on estimated marginal means to better understand any significant effects of these variables on herbivory. To specifically answer question 2b, we added herbivory as a covariate in the Fitness model, i.e. fitness ∼ treatment × species × garden + herbivory + pre-treatment plant size + (1|plot). We compared results of this model to those of the original Fitness Model to determine whether patterns of herbivory mediated the effects of the treatment × species or treatment × species × garden interactions, which would be indicated by a reduction in their *χ^2^* statistics.

Using data from the 2021 field experiment, we re-visited question 1 by running a linear mixed-effects model using a similar formula as above. However, seed number (natural log-transformed) was the response variable representing fitness, and the “garden” variable was removed because the 2021 experiment only occurred in Michigan. Ultimately, we again examined whether a significant treatment × species interaction was supported with this data. If so, we examined species-level tolerance to dicamba drift accordingly and compared results with those of the 2019 study to gauge the reproducibility of our previous results.

Lastly, we conducted structural equation modeling (*piecewiseSEM* package; Lefcheck, 2016) to answer question 3 and determine which functional traits, given dicamba exposure, directly and/or indirectly influence plant fitness as estimated from seed production in the 2021 field experiment. Structural equation modeling is a useful way to test direct and indirect effects by linking multiple variables into a single framework as well as a method that allows for simultaneous testing of multiple hypotheses (Grace, 2006; Lefcheck, 2016). We employed a model selection approach (MSA-SEM), which involves conducting goodness-of-fit tests (via Akaike Information Criterion (AIC) scores) of component models based on a fully-saturated model to identify the most appropriate causal network (Garrido et al., 2021). We performed our analyses by pooling all 9 focal weed species (Baucom et al., 2025; Lázaro et al., 2020), and specifically examined the direct and/or indirect effects of dicamba treatment, leaf damage, 3 week plant size, day of first flower, and flower number on seed number, and any potential interrelationships between these traits.

## Results

### Tolerance of dicamba drift varies by species and environmental context

Over the course of the first experiment (in 2019), the environmental conditions varied by garden. From highest to lowest, the mean ± SE daily temperature was 26.9°C ± 0.20 in Tennessee, 20.4°C ± 0.32 in Michigan, and 19.9°C ± 0.31 in Pennsylvania. The mean ± SE daily precipitation was 25.3 mm ± 6.4 in Tennessee, 18.5 mm ± 5.3 in Michigan, and 45.3 mm ± 8.2 in Pennsylvania. Additionally, as expected, there was a sizable drop in survival for plants that were in the full-dose plots (i.e., were treated with 100% of the field application rate of dicamba). While 81% (209/257) and 92% (191/207) of plants in the control and dicamba drift plots, respectively, survived at three weeks post-treatment, only 25% (8/32) of plants survived in the full-dose plots, confirming that the dicamba herbicide product we used was effective at damaging our study species.

In evaluating the fitness of all nine weed species that were treated with dicamba drift or the control solution across the various gardens, we found that dicamba drift exposure did not have a uniform effect on plant fitness as measured using biomass (main effect of treatment on biomass: *χ^2^* = 0.88, df = 1, *p* = 0.35; Appendix S1: Table S3). Rather, the effect of dicamba drift varied significantly by species (species × treatment effect: *χ^2^* = 19.51, df = 8, *p* = 0.012), indicating that tolerance to dicamba was species-dependent in our constructed weed communities. We also found a significant three-way interaction between species, treatment, and garden, indicating that tolerance to dicamba drift can also be environmentally-dependent (species × treatment × garden effect: *χ^2^* = 19.91, df = 11, *p* = 0.047). Specifically, in the Michigan garden, the biomass of four out of the nine species (*Daucus carota*, *Ipomoea lacunosa*, *I. hederacea*, and *Plantago lanceolata*) was significantly (or marginally significantly) reduced by 76-89% when exposed to dicamba drift (Fig. 1, Appendix S1: Table S4, Appendix S1: S5). In contrast, all species were generally more tolerant of dicamba drift exposure in the Pennsylvania and Tennessee gardens, with the exception of *Amaranthus palmeri* in the Pennsylvania garden, which showed a fairly large reduction (92%; Appendix S1: Table S5) in biomass when treated with dicamba drift.

**Figure 1.**
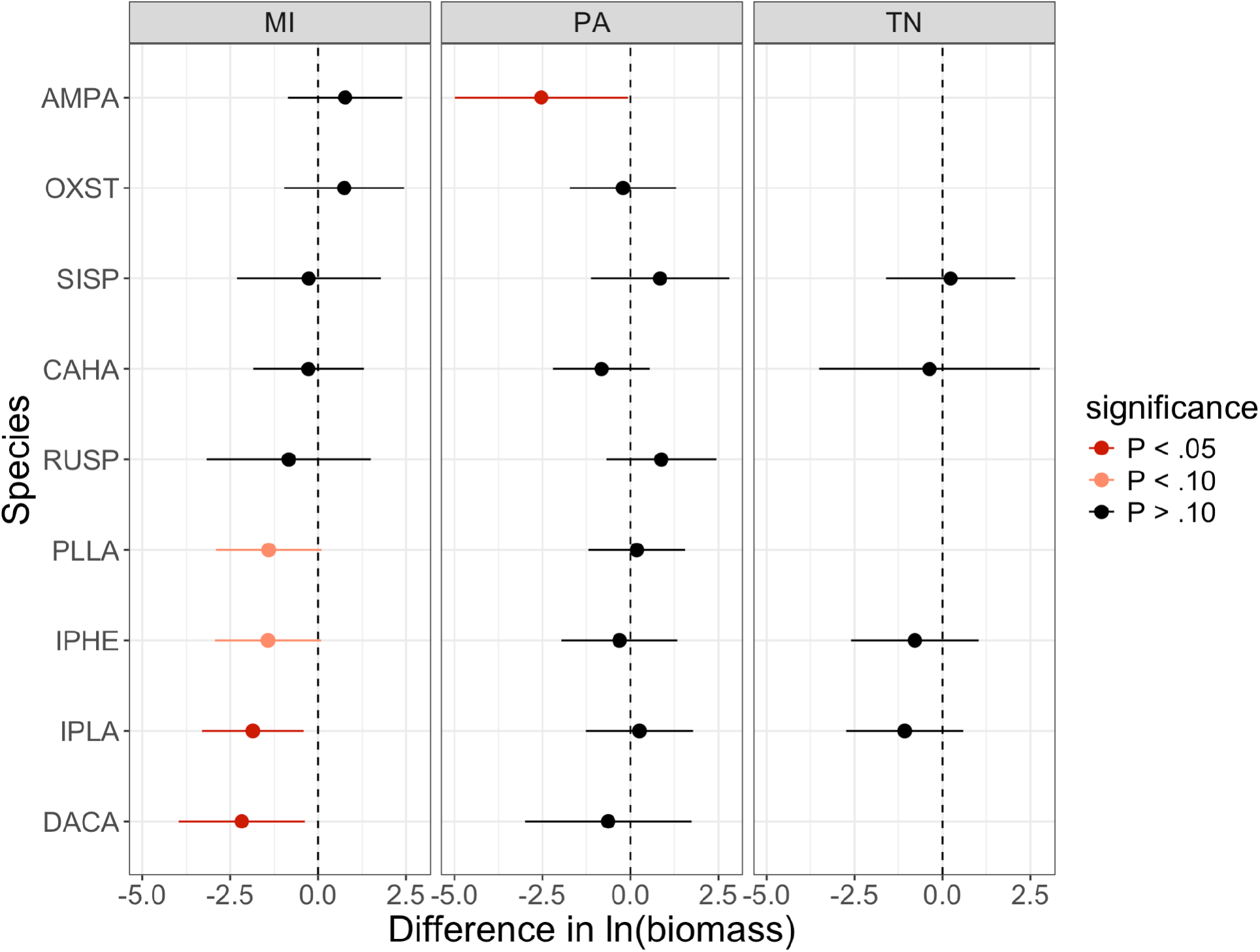
Fitness (biomass) after dicamba drift exposure varies according to weed species and environmental context in a 2019 field study. Points (contrast estimates of estimated marginal means) show the difference in biomass (g) between drift-treated and control-treated plants (biomass_drift_ biomass_control_) of each species (y-axis) at each garden (MI = Michigan, PA = Pennsylvania, TN = Tennessee) in 2019. Error bars show 95% confidence intervals. Colors distinguish between species that were less tolerant to dicamba drift, i.e. their fitness was negatively affected by dicamba drift in a significant (red) or marginally significant (pink) way, and tolerant species whose fitness was unaffected by dicamba drift (black). Contrasts are plotted on the natural log scale because they were natural log-transformed for analysis (see data analysis) and to make comparisons among species easier to appreciate (see Appendix S1: Table S5 for back-transformed values). Species are shown by their four-letter code (Table 1).

In the 2021 experiment, which was conducted only in Michigan and examined plant fitness using seed number, we found similar results regarding the effects of dicamba drift on fitness, but with a few notable differences. In this case, dicamba drift significantly reduced seed set across species by approximately 57% on average (main effect of treatment on seed number: *χ^2^* = 4.59, df = 1, *p* = 0.032; Appendix S1: Table S6). Additionally, *Amaranthus palmeri* was found to be more sensitive to dicamba in this field experiment, showing a 74% reduction in seed number for dicamba-treated plants, whereas in the prior experiment, this species showed no significant change in fitness in the Michigan garden. Congruent however with results from 2019, we found interspecific variation for the effects dicamba drift on fitness (species × treatment effect on seed number: *χ^2^* = 34.69, df = 8, *p* < 0.0001), and we similarly identified that *Daucus carota*, *Ipomoea lacunosa*, and *I. hederacea*, were less tolerant to dicamba, as they produced 80-98% fewer seeds when exposed to the herbicide (Fig. 2; Appendix S1: Table S7, Appendix S1: Table S8). These fitness consequences were slightly more pronounced than what was observed with the biomass data from the first experiment. In comparison, *Oxalis stricta*, *Cardiospermum halicacabum*, and *Sida spinosa* exhibited tolerance to dicamba given that their seed numbers were unaffected by dicamba drift.

**Figure 2.**
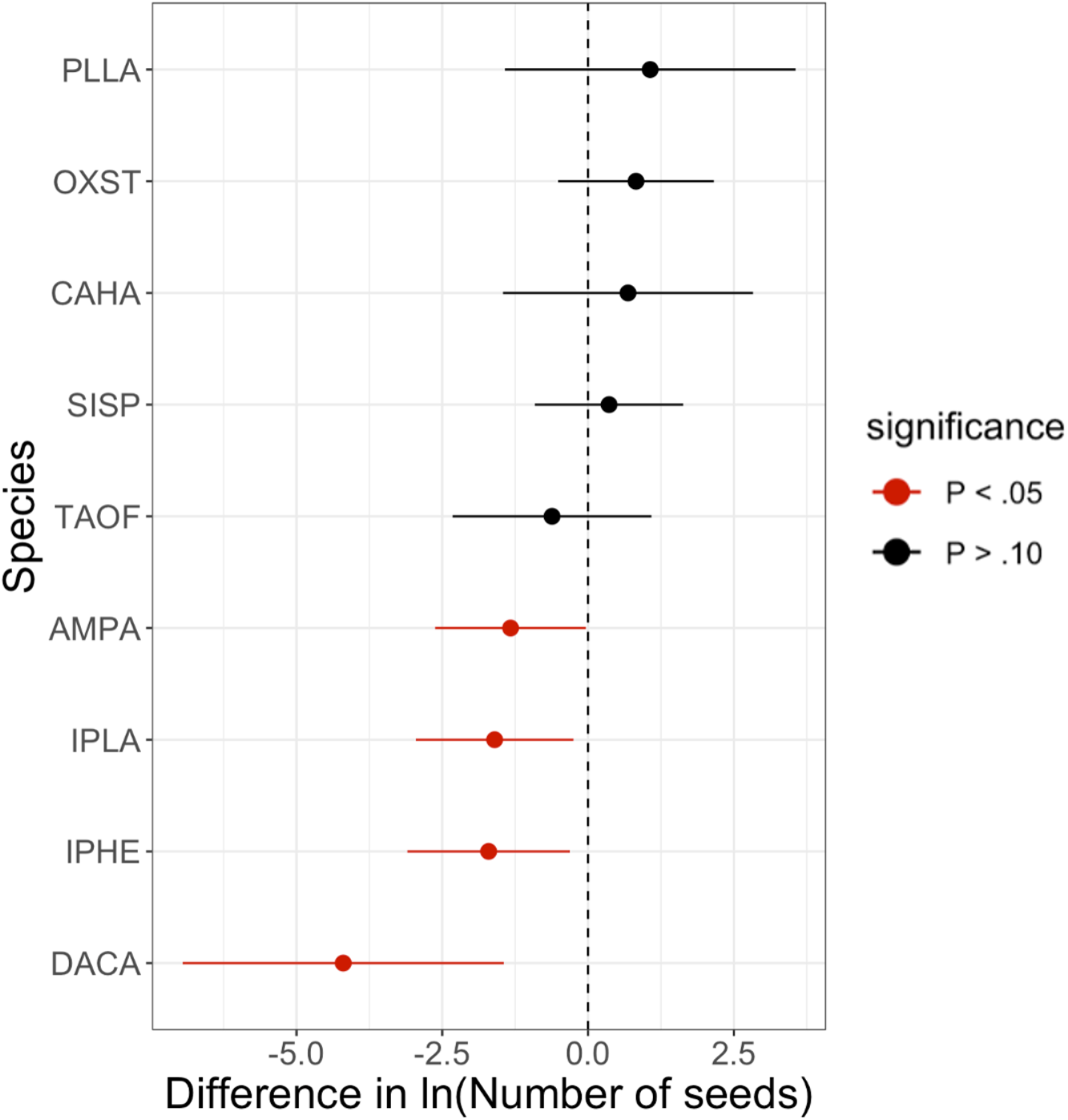
Fitness (seed set) after dicamba drift exposure varies according to weed species in a 2021 field study. Points (contrast estimates of estimated marginal means) show the difference in the number of seeds between drift-treated and control-treated plants (i.e. seed number_drift_ − seed number_control_) of each species (y-axis) in the Michigan garden in 2021. Error bars show 95% confidence intervals. Colors distinguish between species that were less tolerant to dicamba drift, i.e. their fitness was negatively affected by dicamba drift in a significant (red) way, and tolerant species whose fitness was unaffected by dicamba drift (black). Contrasts are plotted on the natural log scale because they were natural log-transformed for analysis (see data analysis) and to make comparisons among species easier to appreciate (see Appendix S1: Table S7 for back-transformed values). Species are shown by their four-letter code (Table 1).

### Herbivory is a significant component of the environmental context and contributes to dicamba drift tolerance

Upon examining the amount of insect herbivory that we observed on experimental plants three weeks after herbicide application, we found that the presence or absence of moderate-severe herbivory was significantly determined by the garden that plants were grown in (*χ^2^*= 14.57, df = 2, *p* = .00069) and their species (*χ^2^*= 46.06, df = 8, *p* <.0001), whereas herbicide treatment did not affect herbivory scores (*χ^2^*= 0.97, df = 1, *p* = 0.33). In particular, plants were significantly more likely to experience moderate-severe herbivory in Pennsylvania (by 18%) and in Tennessee (by 22%) than they were in Michigan (Pennsylvania vs. Michigan: *z*-ratio = −3.50, *p* = 0.0013; Tennessee vs. Michigan: *z*-ratio = −3.035, *p* = 0.0068; Fig. 3A). However, the likelihood of moderate-severe herbivory in Pennsylvania and Tennessee was similar (*z*-ratio = −0.41, *p* = 0.91). In addition, among the species investigated, *Plantago lanceolata*, *Rumex crispus*, *Ipomoea lacunosa,* and *Amaranthus palmeri*, were more likely to be severely damaged by herbivores than the other species, on average (Fig. 3B).

**Figure 3.**
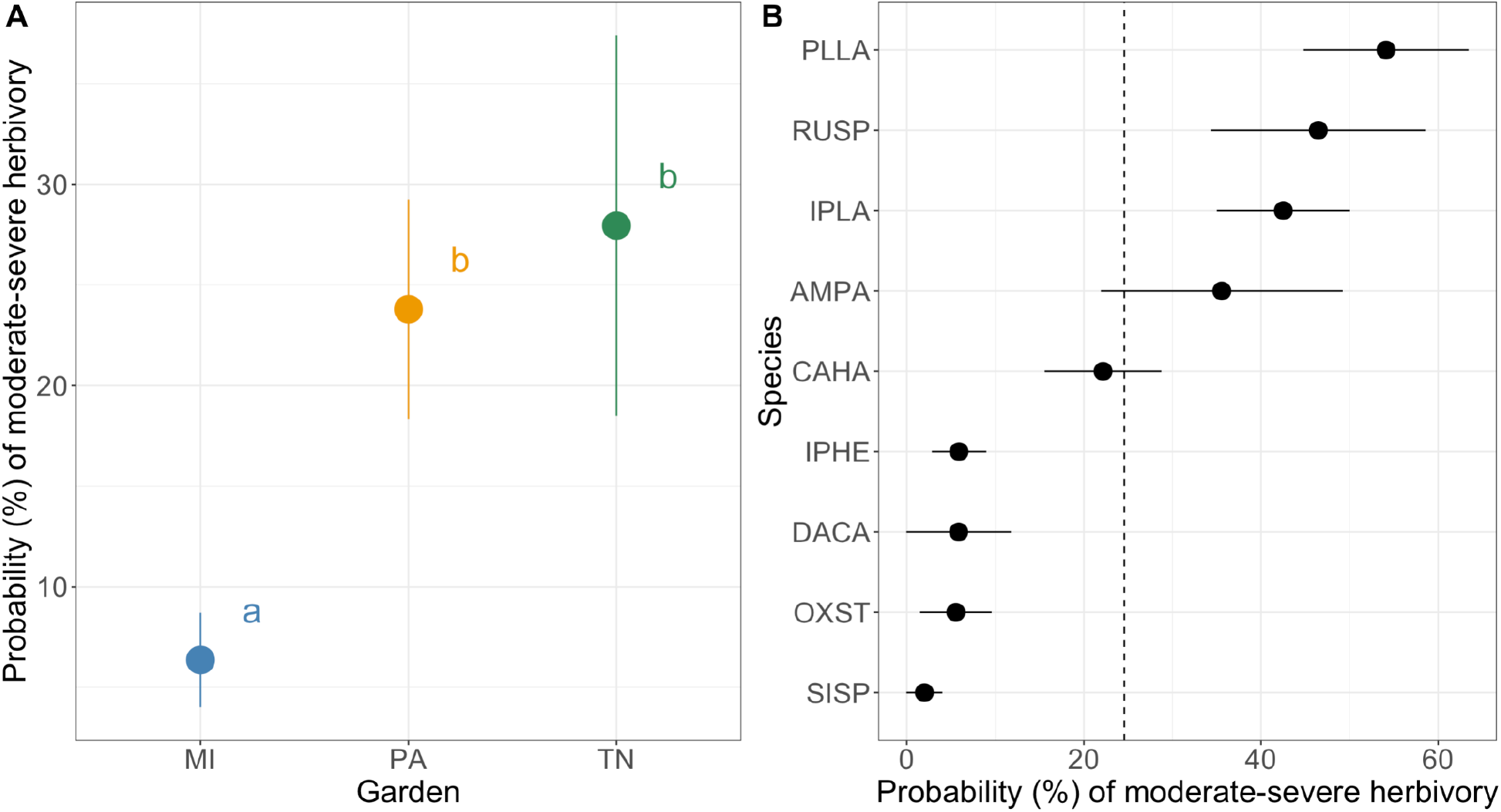
The environmental context and plant species independently affect the probability that plants will experience moderate-severe herbivory. Points show the estimated marginal mean probability that plants will experience moderate-severe herbivory according to: A) the garden they were grown in (MI = Michigan, PA= Pennsylvania, TN = Tennessee), averaged across species, or B) their species, averaged across gardens. In panel A, different letters denote significantly different means between gardens. In panel B, the dotted line indicates the average probability of moderate-severe herbivory across all species, and species are shown by their four-letter code (Table 1). Error bars depict SE.

When we included herbivory as a covariate in our original Fitness Model, we found that, as expected, insect herbivory had a significant effect on plant biomass (*χ^2^* = 6.06, df = 2, *p* = 0.014). Plants that had experienced moderate to severe levels of damage from herbivory were on average 51% smaller than plants that had experienced minimal damage from herbivory. Interestingly, including this term also resulted in a reduced species x treatment x garden effect on biomass that became nonsignificant (*χ^2^* = 13.48, df = 11, *p* = 0.263; Appendix S1: Table S9), as did the species x treatment effect (*χ^2^* = 13.18, df = 8, *p* = 0.106; Appendix S1: Table S9). Altogether, the results from these analyses suggest that the species in our study experienced different levels of herbivory that affected their biomass and were intensified by the environment. Additionally, the fact that herbicide treatment effects on plant biomass were weakened when accounting for the degree of herbivory that occurred post-treatment indicates that herbivory likely played a role in influencing how these species were affected by dicamba drift across the different gardens.

### Early growth and flowering traits underlie the effect of dicamba drift on the fitness of weed species

We determined the best-fit structural equation model that could describe potential relationships between dicamba exposure, plant traits (leaf damage, 3wk plant size, day of first flower, and flower number), and fitness using a model selection approach (Fig. 4; see Appendix S1: Table S10 for model selection results). This model met the goodness-of-fit threshold (Fisher’s *C* = 1.89, *p* = 0.76), indicating that it was a valid representation of the underlying data and was not missing any key pathways (Lefcheck, 2016). The model uncovered four distinct pathways by which dicamba drift exposure indirectly affected plant fitness as determined by seed number in the 2021 field experiment, with a negative total effect of −0.36 (Table 2).

**Figure 4.**
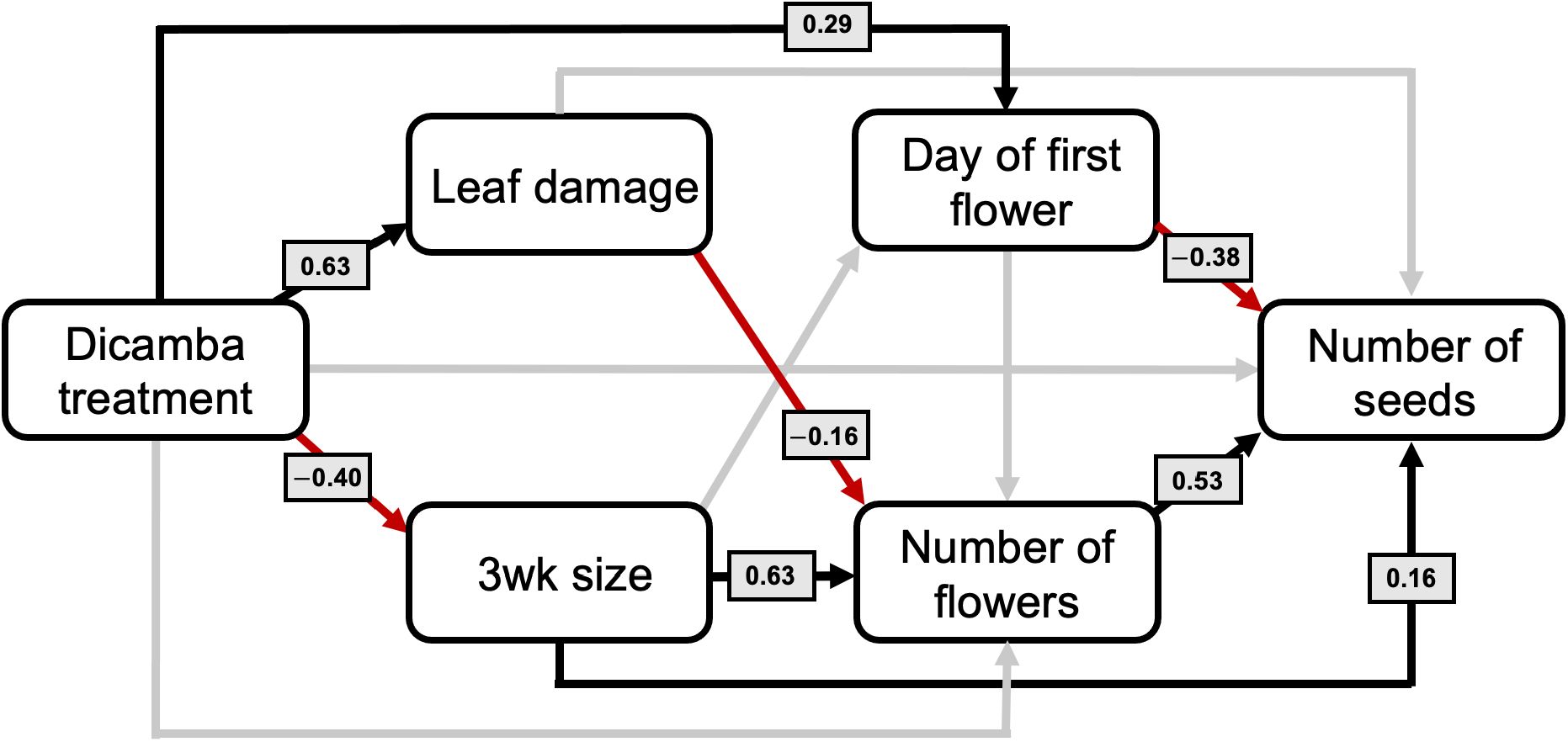
Structural equation modeling identifies traits underlying seed production following exposure to dicamba drift across weedy plant species. Potential relationships between dicamba drift exposure (dicamba treatment), vegetative traits (3wk plant size, leaf damage), floral traits (day of first flower, number of flowers), and seed production (number of seeds) are presented from the best-fit model structural equation model (Appendix S1: Table S10). Red lines indicate significant negative relationships, whereas black lines depict significant positive relationships. Nonsignificant paths are shown in grey.

**Table 2.** The pathways describing the indirect effects of dicamba exposure on seed production.

| Pathway | Description | Total effect on number of seeds |
| --- | --- | --- |
| 1 | Dicamba treatment → 3wk size → Seed production | -0.064 |
| 2 | Dicamba treatment → 3wk size → Number of flowers → Seed production | -0.13 |
| 3 | Dicamba treatment → Leaf damage → Number of flowers → Seed production | -0.053 |
| 4 | Dicamba treatment → Day of first flower → Seed production | -0.11 |
|  | **Grand total** | **-0.36** |

Notes: The total effect of each pathway was determined by multiplying the standardized path coefficients across the pathway. A grand total effect for each metric of seed production (number of seeds) was calculated by summing across pathways. “NA” indicates that the pathway was not substantiated by significant path coefficients throughout the pathway.

Specifically, dicamba exposure reduced plant size in the short term, which reduced seed number directly by −0.064 (Pathway 1) as well as indirectly through effects on flower number (Pathway 2). Because short-term plant size had a large positive effect of 0.63 on flower number, the dicamba drift treatment ultimately reduced seed number indirectly by −0.13 *via* this pathway. Additionally, greater leaf damage caused by dicamba exposure directly led to decreased flower numbers (by −0.16) and thereby indirectly decreased seed number by −0.053 (Pathway 3). Finally, while dicamba drift did not influence flower number directly, it directly delayed flowering onset by 0.29 (positive values for day of first flower indicate later flowering; Fig. 4), and delayed flowering resulted in direct reductions in seed number by −0.38, such that the total indirect effect of dicamba on seed number through this pathway (Pathway 4) was −0.11. Thus, these results suggest that short-term changes in vegetative traits within the first few weeks of dicamba exposure (which result in decreases in flower production) and direct delays in flowering phenology are especially important mechanisms by which dicamba drift can reduce the fitness of various species of common weeds, thus influencing their ability to tolerate the herbicide.

## Discussion

Through exploring the effects of sublethal drift exposures from the frequently-used synthetic auxin herbicide dicamba on constructed plant communities in the field, we discovered that the effects of dicamba drift are both species and context dependent: species varied in their tolerance to dicamba exposure and variation in tolerance among species was also garden-dependent. We also found that herbivory played a role in determining whether species showed variation in tolerance to dicamba drift across three different agro-eco environments. Further, our use of structural equation modeling allowed us to clarify key functional pathways by which dicamba drift exerts its effects on plant fitness–namely, dicamba drift reduces seed production by disrupting short-term growth and flower production. Here, we discuss the potential causes of these newly observed patterns and their ecological implications for present-day agroecosystems.

### Unpacking interspecific variation in dicamba drift tolerance among weedy plants under field conditions

Our monitoring of nine common crop-associated weed species across three different environmental contexts in the central region of the United States found that species varied considerably in their ability to tolerate dicamba drift exposure. Whereas some species were able to maintain fitness despite having been exposed to the herbicide, others showed severe (up to 90%) reductions in shoot biomass by the end of the growing season. This provides support to previous studies (Baucom et al., 2025; Iriart et al., 2022; Olszyk et al., 2017), showing that even very low-dose exposures to dicamba can weaken certain members of weedy plant communities, which could influence the shift of these communities to favor dicamba-tolerant species (e.g. *Oxalis stricta*, *Rumex crispus*, and *Cardiospermum halicacabum*; Fig. 1). Yet, it is notable that two field studies (here, Baucom et al., 2025) found less appreciable effects of dicamba than a greenhouse study by Iriart et al. (2022). Specifically, we did not find evidence for overcompensation, and a smaller proportion of the species investigated exhibited statistically significant reductions in biomass attributable to dicamba drift. Harsher conditions in the field compared to the greenhouse could explain this result, as less favorable environments limit growth (Conner et al., 2003). This hypothesis could likewise be applied to explain the differences in interspecific variation we observed among the three field environments (i.e. gardens), as discussed below.

Interestingly, while our analysis of seed number data in the 2021 field experiment generally confirmed our findings using biomass to estimate fitness in the 2019 experiment, we found that the effect of dicamba drift on seed number was greater than that of biomass. Shoot biomass is often used as a proxy for fitness because it is relatively accessible and often correlates with fecundity-related traits (reviewed in Younginger et al., 2017), but there are limitations to using this approach. Different environmental conditions, including the availability of resources (e.g. water, nutrients, pollinators and other mutualists) can result in discrepancies between biomass and reproductive responses (Burkle & Irwin, 2010). Given that our study species (like many weedy species) are predominantly insect-pollinated (Table 1), it is possible that differences in pollinator accessibility mediated the difference in the overall treatment effect. Namely, a reduction in pollinator visitation to drift-exposed plant communities as suggested by Baucom et al. (2025) and Bohnenblust et al. (2016) could have resulted in more pollen-limited plants with reduced seed set (Ashman et al., 2004). That said, we still found that the patterns which characterized species-level variation in tolerance using biomass as a fitness estimate in 2019 generally recapitulated those using seed number in the experiment performed in 2021 in Michigan (Fig. 1, Fig. 2).

### Environment-dependent effects of dicamba drift on weed species

While interspecific variation in dicamba response was clearly seen in the Michigan garden in our 2019 study, it was less evident in the Pennsylvania and Tennessee gardens. This may be because the Michigan garden was more favorable, with fewer added stressors than the other two gardens. For example, although we did not observe an increase in herbivory damage caused by dicamba directly as in previous work on a smaller number of weedy plant species (Johnson et al., 2019; Johnson et al., 2022), we found that changes in herbivory degree following herbicide application did account for a considerable amount of the variation in biomass observed across species, treatments, and gardens (Appendix S1: Table S6). Thus, it is likely that herbivory mediates the strength by which weed species respond to dicamba drift in varying contexts. In particular, plants in the Pennsylvania and Tennessee gardens experienced greater herbivory pressure than those in the Michigan garden (Fig. 3; Appendix S1: Table S6). Consequently, the effect of dicamba on plants in Michigan was likely more isolated, such that control plants were able to reach a higher fitness, resulting in greater fitness differences between control and dicamba-treated plants. This outcome was probably most applicable for species that tended to be more heavily herbivorized, such as *Ipomoea lacunosa* and *Plantago lanceolata* (Fig. 3B), whose sensitivity to dicamba drift in the Michigan garden may have been more apparent because of the lack of additional pressure from herbivores. It is important to note, however, that while we roughly estimated herbivory damage, future research should consider using a finer scale (e.g. by quantifying missing tissue and mode of damage; Zhang & Baucom, 2024) to draw greater insights into the relationship between herbivory and response to herbicide stress.

The climatic trends which occurred over the course of the study also support the hypothesis that interspecific variation in herbicide tolerance is more appreciable in the absence of additional environmental stressors: while the Michigan garden had the mildest climate over the course of the growing season (daily temperature: 20°C, precipitation: 19 mm), the Tennessee garden experienced higher summer temperatures (daily average: 27°C), and the Pennsylvania garden experienced higher precipitation (daily average 45 mm). However, one exception to this pattern was the tolerance results of *Amaranthus palmeri*, a species that exhibited greater differences between treated and untreated plants in Pennsylvania than in Michigan. Although *A. palmeri* is typically associated with arid conditions, it is also known for being highly adaptable (it often top lists of problematic agricultural weeds; Chahal et al., 2018), and some research speculates that it may show increased growth relative to other plants in instances of high precipitation or water-logging (Zhang et al., 2022). Therefore, it is possible that while control-treated *A. palmeri* plants could withstand the wetter conditions in Pennsylvania, disruptions in water usage caused by synthetic auxin function (Suliman et al., 2025) may have impeded growth for drift-treated *A. palmeri*. These findings warrant further investigation into potential interactions between climatic variables and herbicidal efficacy at sublethal doses.

### Linking plant functional traits to dicamba drift tolerance

Our structural equation model allowed us to more clearly understand the process driving these fitness differences between weedy plants that are affected or unaffected by herbicide drift. Through it, we demonstrated that leaf damage and changes in plant size caused by dicamba in the short term have significant downstream consequences for flower production and fitness in the long term (Fig. 4). This both confirmed patterns that had been predicted previously (Iriart et al., 2020; Iriart et al., 2022) and elucidated nuanced relationships. For example, although leaf damage is the most commonly reported form of injury to nontarget plants exposed to dicamba drift (Buol et al., 2019; Johnson et al., 2023; Kniss et al., 2018), the phenotypic changes caused by dicamba treatment that most strongly related to fitness across the weed species investigated were less conspicuous—namely, changes in plant size resulted in reductions in flower number that reduced fitness, and changes in the initiation of flowering decreased fitness (Fig. 4; Table 2). Based on key findings from Baucom et al. (2025) that investigated the same set of plants, it is likely that reductions and delays in flower production instigated by dicamba drift resulted in significant decreases in pollinator visitation rates, which in turn would have facilitated plant reproduction (Ashman et al., 2004).

Taken together, these results indicate that the weedy species that are most likely to dominate communities that commonly encounter dicamba drift are probably species that can mitigate changes in early growth and flowering post-dicamba exposure. Concomitantly, they suggest that herbicide drift-tolerant species of weeds are not likely to experience a fitness cost in allocating resources towards growth maintenance in the short term. An appropriate next step of this research would be to closely examine a select set of dicamba drift-tolerant and undercompensating weed species to address whether there are any species-specific differences in how these functional traits relate to dicamba drift tolerance. Additionally, extending our experimental set-up over multiple years would shed light on whether shifts in weed communities occur as predicted, i.e. whether tolerant or overcompensating species reach greater population sizes than undercompensating species in drift-exposed landscapes over time.

## Conclusion

Overall, our study supports previous findings that low-dose exposures to an auxin-mimicking herbicide differentially affects plant species via changes in vegetative size and flower production. It also provides new insights about the context-dependent nature of the effects of this form of anthropogenic stress on “weedy” plants, and bridges the gap between observations of species differences in performance in the greenhouse and those in natural environments. To further build upon this knowledge base, future studies should consider other variables that could explain the relationship between the environment and plant fitness outcomes under herbicide stress. For example, other critical factors that may mediate synthetic auxin herbicide action, in addition to the ones discussed above, include soil type and plant-microbe interactions in the rhizosphere (Chowdhury et al., 2025; Iriart et al., 2024; Zhu et al., 2015). Looking forward, the use of synthetic auxin herbicides is growing on a broad scale (Garcia et al., 2025), as is public concern surrounding biodiversity loss, especially among native pollinators (Cornelisse et al., 2025). As such, understanding the unintentional consequences of agrochemical movement onto weedy plant communities–upon which pollinators and other wildlife rely–remains a paramount issue for both basic and applied ecological research.

## Acknowledgements

Our research was supported by USDA 2017-09529/1016564 to T.-L.A. and R.S.B. V.I. was additionally supported by the University of Pittsburgh Dietrich School of Arts and Sciences First-Year and K. Leroy Irvis Fellowships. We thank A. Blanco and E. O’Neill for their assistance in data collection, and L. Follweiler for applying herbicide treatments at the Pennsylvania garden. We thank Mike Palmer and Jeremy Moghtader and the Matthaei Botanical Gardens for assistance with field preparation and maintenance at the Michigan garden and the Long Family for the use of their land for the Tennessee garden.

## Author Contributions

V.I., R.S.B., and T.L.-A. conceived and designed the study. V.I., A.S.B., T.N., S.C., and M.A. performed the study under supervision by T.L.-A and R.S.B. V.I. analysed the data, and wrote the first draft of the manuscript, wherein R.S.B. and T.L.-A. contributed revisions.

