## Appendix S1 for "Tolerance to herbicide drift in “weedy” species is context-dependent and driven by early vegetative growth"

Supporting information for the manuscript: “Tolerance to herbicide drift in ‘weedy’ species is context-dependent and driven by early vegetative growth”

Veronica Iriart, Tia-Lynn Ashman, Matthew R. Armstrong, Sara Colom, Toshiro Newsum, Anah Soble Botran, and Regina S. Baucom

**Table S1. Information on all plant species planted in the 2019 field experiment and their seed sources.** See Table S2 for location information of all seed collection sites.

| **Species** | **Family** | **Seed collection sites** |
|---|---|---|
| *Acalypha ostryifolia* | Euphorbiacea | 18, 27, 25 |
| *Amaranthus palmeri* | Amaranthacea | 15, 22, DG07 |
| *Asclepias syriaca* | Aslepias | 17 |
| *Cardiospermum halicacabum* | Sapindaceae | 17, HJ03 |
| *Chamaecrista fasciculata* | Fabaceae | 22, 26A |
| *Cirsium arvense* | Asteraceae | HJ04 |
| *Conyza canadensis* | Asteraceae | 24 |
| *Daucus carota* | Apiaceae | 33A, 24 |
| *Desmanthus illinoensis* | Fabaceae | HJ01, HJ04 |
| *Erigeron annuus* | Asteraceae | 22, 24 |
| *Euphorbia maculata* | Euphorbiaceae | 28, 29A, 31 |
| *Ipomoea lacunosa* | Concolvulacea | DG07, 11, HJ03 |
| *Ipomoea purpurea* | Concolvulacea | DG08, 27A |
| *Lepidium virginicum* | Brassicaceae | 14, 22, 29A |
| *Medicago lupulina* | Fabaceae | 29A, 30, 27 |
| *Mollugo verticillata* | Mollugoinacea | 22, 24, DG08 |
| *Oxalis stricta* | Oxalidaceae | 30A, 24, 32A |
| *Persicaria spp.* | Polygonaceae | 13, 14, 28 |
| *Plantago lanceolata* | Plantaginaceae | 13, 29A, 33A |
| *Rumex spp.* | Polygonaceae | 23, 24 |
| *Senna obtusifolia* | Fabaceae | HJ01 |
| *Sida spinosa* | Malvaceae | 21, 31, HJ01 |
| *Solidago canadensis* | Asteracea | 24, 26, 36A |
| *Taraxacum officinale* | Asteraceae | HJ01, 19, 24 |
| *Trifolium pretense* | Fabaceae | 27, 33A, 29A |
| *Trifolium repens* | Fabaceae | DG08, 12, 19 |
| *Veronia fasciculata* | Asteraceae | 32A |

**Table S2. Location information of sites visited to collect seeds in the 2019 field experiment.**

| Site | Latitude | Longitude | County, State in USA |
|---|---|---|---|
| 11 | 36.5976 | -86.8511 | Robertson County, TN |
| 13 | 36.7988 | -86.8762 | Logan County, KY |
| 14 | 37.0655 | -86.6151 | Warren County, KY |
| 15 | 37.1281 | -86.6596 | Butler County, KY |
| 17 | 36.9771 | -86.7835 | Logan County, KY |
| 19 | 36.8153 | -87.1768 | Todd County, KY |
| 20 | 36.8494 | -87.3016 | Christian County, KY |
| 21 | 36.854 | -87.5514 | Christian County, KY |
| 22 | 36.7271 | -88.1781 | Calloway County, KY |
| 23 | 36.6906 | -88.189 | Calloway County, KY |
| 24 | 36.6614 | -88.2168 | Calloway County, KY |
| 25 | 36.5696 | -88.2472 | Calloway County, KY |
| 26 | 36.5347 | -87.5709 | Christian County, KY |
| 26A | 36.5346 | -87.5658 | Christian County, KY |
| 27 | 36.8133 | -87.4098 | Christian County, KY |
| 27A | 36.8257 | -87.477 | Christian County, KY |
| 28 | 36.8993 | -87.5339 | Christian County, KY |
| 29A | 36.7995 | -87.3839 | Christian County, KY |
| 30 | 36.7929 | -87.3796 | Christian County, KY |
| 30A | 36.7929 | -87.3796 | Christian County, KY |
| 31 | 36.7014 | -87.0415 | Logan County, KY |
| 32A | 36.6409 | -87.1576 | Montgomery County, TN |
| 33A | 36.44 | -86.8106 | Robertson County, TN |
| DG07 | 36.6233 | -86.8413 | Robertson County, TN |
| DG08 | 36.6319 | -86.8636 | Robertson County, TN |
| HJ01 | 35.751 | -86.6208 | Rutherford County, TN |
| HJ03 | 35.7333 | -86.5902 | Rutherford County, TN |
| HJ04 | 35.7333 | -86.5902 | Rutherford County, TN |
| JB06 | 35.8729 | -86.4745 | Rutherford County, TN |

**Table S3. Results of type III sum of squares ANOVA analyzing biomass in the 2019 field study**. Abbreviations: df = degrees of freedom. Significant *p* values are bolded.

| **Fixed Effect** | $\chi^2$ | **df** | ***p*** |
|---|---|---|---|
| Treatment (TRT) | 0.88 | 1 | 0.349 |
| Species (SPP) | 59.28 | 8 | **<.0001** |
| Garden | 18.75 | 2 | **<.0001** |
| Pre-treatment plant size | 60.52 | 1 | **<.0001** |
| TRT × SPP | 19.51 | 8 | **0.012** |
| TRT × Garden | 5.25 | 2 | 0.073 |
| SPP × Garden | 19.70 | 11 | **0.0496** |
| TRT × SPP × Garden | 19.66 | 11 | **0.047** |

**Table S4. Contrast estimates of the difference in biomass between plants treated with dicamba drift vs. control according to species and garden.** Differences were calculated by subtracting mean biomass in drift by mean biomass in control, using estimated marginal means. Estimates and confidence levels are on the natural log scale. Abbreviations: df = degrees of freedom, LCL = lower confidence level, UCL = upper confidence level. Significant *p* values are bolded.

| Species | Garden | Estimate | df | LCL | UCL | *t* ratio | *p* |
|---|---|---|---|---|---|---|---|
| AMPA | MI | 0.768 | 98 | -0.86 | 2.40 | -0.94 | 0.352 |
| CAHA | MI | -0.278 | 94 | -1.85 | 1.29 | 0.35 | 0.726 |
| DACA | MI | -2.173 | 138 | -3.97 | -0.38 | 2.40 | **0.018** |
| IPHE | MI | -1.426 | 80 | -2.94 | 0.08 | 1.88 | 0.064 |
| IPLA | MI | -1.860 | 69 | -3.31 | -0.41 | 2.56 | **0.013** |
| OXST | MI | 0.743 | 117 | -0.96 | 2.45 | -0.86 | 0.390 |
| PLLA | MI | -1.407 | 78 | -2.91 | 0.09 | 1.87 | 0.066 |
| RUSP | MI | -0.837 | 233 | -3.17 | 1.50 | 0.71 | 0.481 |
| SISP | MI | -0.268 | 185 | -2.31 | 1.78 | 0.26 | 0.796 |
| AMPA | PA | -2.531 | 249 | -4.99 | -0.07 | 2.03 | **0.044** |
| CAHA | PA | -0.821 | 60 | -2.20 | 0.56 | 1.19 | 0.239 |
| DACA | PA | -0.628 | 241 | -3.00 | 1.74 | 0.52 | 0.602 |
| IPHE | PA | -0.308 | 108 | -1.95 | 1.34 | 0.37 | 0.711 |
| IPLA | PA | 0.261 | 84 | -1.26 | 1.79 | -0.34 | 0.735 |
| OXST | PA | -0.208 | 82 | -1.72 | 1.30 | 0.27 | 0.785 |
| PLLA | PA | 0.184 | 60 | -1.19 | 1.56 | -0.27 | 0.790 |
| RUSP | PA | 0.882 | 93 | -0.68 | 2.44 | -1.12 | 0.265 |
| SISP | PA | 0.846 | 177 | -1.12 | 2.82 | -0.85 | 0.398 |
| CAHA | TN | -0.367 | 238 | -3.51 | 2.78 | 0.23 | 0.818 |
| IPHE | TN | -0.785 | 68 | -2.60 | 1.03 | 0.86 | 0.391 |
| IPLA | TN | -1.072 | 47 | -2.73 | 0.59 | 1.30 | 0.200 |
| SISP | TN | 0.232 | 84 | -1.60 | 2.07 | -0.25 | 0.802 |

**Table S5. Back-transformed estimated marginal means and contrast estimates of plant biomass according to species and garden.** "Control emmean" and "Control SE" are the estimated marginal mean biomass and its standard error, respectively, for plants treated with the control. "Drift emmean" and "Drift SE" are the estimated marginal mean biomass and its standard error, respectively, for plants treated with dicamba drift. "Contrast estimate", "Contrast LCL", and "Contrast UCL" are the difference in the estimated marginal means between control and drift-treated plants (Drift − Control), the lower confidence limit of this difference, and the upper confidence limit of this difference, respectively. Values were back-transformed to the original scale of measurement (g).

| Species | Garden | Control emmean | Control SE | Drift emmean | Drift SE | Contrast estimate | Contrast LCL | Contrast UCL |
|---|---|---|---|---|---|---|---|---|
| AMPA | MI | 34.37 | 18.54 | 74.07 | 46.10 | 39.70 | -134.39 | 213.80 |
| CAHA | MI | 23.10 | 12.80 | 17.48 | 10.01 | -5.61 | -61.90 | 50.67 |
| DACA | MI | 5.32 | 3.65 | 0.61 | 0.36 | -4.71 | -17.38 | 7.95 |
| IPHE | MI | 68.89 | 34.25 | 16.55 | 9.51 | -52.33 | -178.28 | 73.61 |
| IPLA | MI | 21.17 | 10.40 | 3.30 | 1.77 | -17.87 | -55.27 | 19.53 |
| OXST | MI | 1.05 | 0.67 | 2.21 | 1.30 | 1.16 | -3.89 | 6.20 |
| PLLA | MI | 9.43 | 4.83 | 2.31 | 1.28 | -7.12 | -24.72 | 10.48 |
| RUSP | MI | 2.48 | 2.07 | 1.07 | 0.91 | -1.41 | -9.12 | 6.30 |
| SISP | MI | 5.27 | 4.48 | 4.03 | 2.42 | -1.24 | -18.78 | 16.31 |
| AMPA | PA | 0.64 | 0.48 | 0.05 | 0.05 | -0.59 | -2.25 | 1.07 |
| CAHA | PA | 6.58 | 3.14 | 2.90 | 1.45 | -3.69 | -16.01 | 8.64 |
| DACA | PA | 0.96 | 0.95 | 0.51 | 0.35 | -0.45 | -3.93 | 3.03 |
| IPHE | PA | 8.28 | 4.37 | 6.08 | 3.90 | -2.20 | -22.70 | 18.31 |
| IPLA | PA | 10.21 | 5.45 | 13.24 | 7.30 | 3.04 | -28.85 | 34.93 |
| OXST | PA | 0.43 | 0.22 | 0.35 | 0.20 | -0.08 | -1.13 | 0.96 |
| PLLA | PA | 1.42 | 0.71 | 1.71 | 0.82 | 0.29 | -3.56 | 4.13 |
| RUSP | PA | 0.37 | 0.23 | 0.90 | 0.47 | 0.53 | -1.28 | 2.34 |
| SISP | PA | 1.21 | 0.89 | 2.83 | 1.92 | 1.61 | -5.64 | 8.87 |
| CAHA | TN | 1.36 | 1.93 | 0.94 | 0.68 | -0.42 | -7.31 | 6.48 |
| IPHE | TN | 2.11 | 1.43 | 0.96 | 0.58 | -1.15 | -6.52 | 4.23 |

| | | | | | | | | |
|---|---|---|---|---|---|---|---|---|
| IPLA | TN | 1.93 | 1.21 | 0.66 | 0.40 | -1.27 | -5.41 | 2.87 |
| SISP | TN | 1.92 | 1.33 | 2.42 | 1.47 | 0.50 | -6.61 | 7.61 |

**Table S6. Results of type III sum of squares ANOVA analyzing seed number in the 2021 field study**. Abbreviations: df = degrees of freedom. Significant *p* values are bolded.

| Fixed Effect | $\chi^2$ | df | *p* |
|---|---|---|---|
| Treatment (TRT) | 4.59 | 1 | **0.032** |
| Species (SPP) | 92.36 | 8 | **<.0001** |
| Pre-treatment plant size | 42.35 | 1 | **<.0001** |
| TRT × SPP | 34.69 | 8 | **<.0001** |

**Table S7. Contrast estimates of the difference in seed number between plants treated with dicamba drift vs. control according to species and garden.** Differences were calculated by subtracting seed number in drift by seed number in control, using estimated marginal means for seed number. Estimates and confidence levels are on the natural log scale. Abbreviations: df = degrees of freedom, LCL = lower confidence level. UCL = upper confidence level. Significant *p* values are in bold.

| Species | Estimate | df | LCL | UCL | *t* ratio | *P* |
|---|---|---|---|---|---|---|
| AMPA | -1.33 | 21 | -2.62 | -0.04 | 2.14 | **0.044** |
| CAHA | 0.68 | 113 | -1.46 | 2.83 | -0.63 | 0.528 |
| DACA | -4.20 | 166 | -6.95 | -1.44 | 3.01 | **0.003** |
| IPHE | -1.71 | 30 | -3.10 | -0.31 | 2.50 | 0.018 |
| IPLA | -1.60 | 26 | -2.95 | -0.25 | 2.44 | **0.022** |
| OXST | 0.82 | 25 | -0.51 | 2.16 | -1.27 | 0.217 |
| PLLA | 1.06 | 150 | -1.43 | 3.56 | -0.84 | 0.400 |
| SISP | 0.36 | 20 | -0.91 | 1.63 | -0.59 | 0.562 |
| TAOF | -0.62 | 65 | -2.32 | 1.09 | 0.72 | 0.472 |

**Table S8. Back-transformed estimated marginal means and contrast estimates of seed set according to species and garden.** "Control emmean" and "Control SE" are the estimated marginal mean number of seeds and its standard error, respectively, for plants treated with the control treatment. "Drift emmean" and "Drift SE" are the estimated marginal mean number of seeds and its standard error, respectively, for plants treated with the dicamba drift treatment. "Contrast estimate", "Contrast LCL", and "Contrast UCL" are the difference in the estimated marginal means between control and drift-treated plants (Drift − Control), the lower confidence limit of this difference, and the upper confidence limit of this difference, respectively. Values were back-transformed to the original scale of measurement (seed counts).

| Species | Emmean in Control | SE in Control | Emmean in Drift | SE in Drift | Contrast estimate | Contrast LCL | Contrast UCL |
|---|---|---|---|---|---|---|---|
| AMPA | 6878 | 3048 | 1820 | 802 | -5058 | -16489 | 6373 |
| CAHA | 274 | 167 | 543 | 484 | 269 | -1408 | 1946 |
| DACA | 8239 | 7433 | 124 | 132 | -8115 | -31708 | 15478 |
| IPHE | 253 | 121 | 46 | 22 | -207 | -636 | 222 |
| IPLA | 251 | 128 | 51 | 21 | -200 | -658 | 257 |
| OXST | 1902 | 911 | 4327 | 1939 | 2425 | -4983 | 9833 |
| PLLA | 73 | 66 | 212 | 186 | 139 | -489 | 766 |
| SISP | 592 | 265 | 849 | 353 | 257 | -1339 | 1853 |
| TAOF | 343 | 205 | 185 | 113 | -158 | -925 | 608 |

**Table S9. Results of type III sum of squares ANOVA analyzing biomass in the 2019 field study with the presence/absence of herbivory included as a covariate**. Abbreviations: df = degrees of freedom. Significant *p* values are bolded.

| **Fixed Effect** | $\chi^2$ | **df** | ***p*** |
|---|---|---|---|
| Treatment (TRT) | 0.01 | 1 | 0.93 |
| Species (SPP) | 68.63 | 8 | **<.0001** |
| Garden | 20.45 | 2 | **<.0001** |
| Pre-treatment plant size | 61.27 | 1 | **<.0001** |
| Herbivory | 6.06 | 1 | **0.014** |
| TRT × SPP | 13.18 | 8 | 0.106 |
| TRT × Garden | 2.68 | 2 | 0.262 |
| SPP × Garden | 23.27 | 11 | **0.016** |
| TRT × SPP × Garden | 13.48 | 11 | 0.263 |

**Table S10. Results of model selection comparing structural equation models (SEMs) for best fit.** SEMs, as depicted in the figures shown in "Model", were constructed to investigate potential relationships between dicamba drift (dicamba treatment), vegetative traits (3wk plant size, leaf damage), floral traits (day of first flower, number of flowers), and seed production (number of seeds). "AIC" shows the Akaike Information Criterion (AIC) score for the constructed model and "*Δ*AIC" shows the difference in the AIC score between the constructed model and the one with the lowest AIC score, i.e. the highest ranking model that best-fit the data.

| Rank | Model | AIC | *Δ*AIC |
|---|---|---|---|
| 1 | 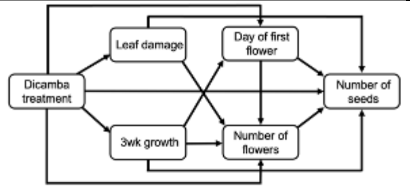 | 57.89 | 0.00 |
| 2 | 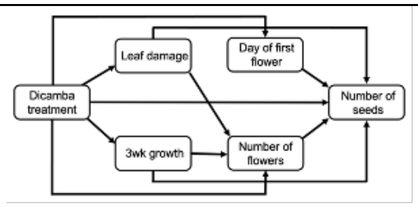 | 58.34 | 0.45 |
| 3 | 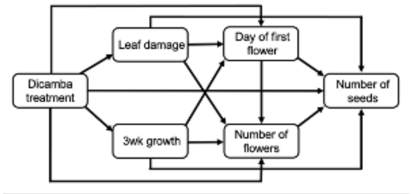 | 59.09 | 1.20 |
| 4 | 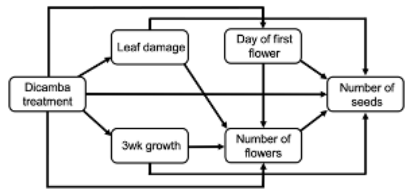 | 59.96 | 2.07 |
| 5 | 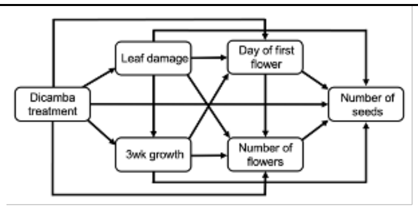 | 60.00 | 2.11 |
| 6 | 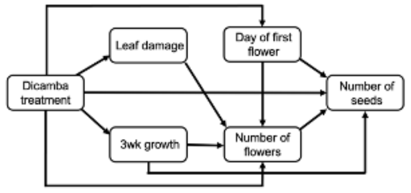 | 60.87 | 2.98 |

| | | | |
|---|---|---|---|
| 7 | 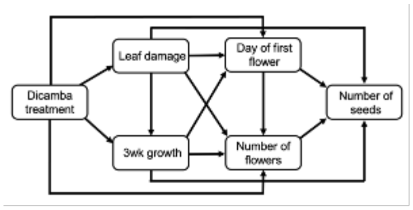 | 61.51 | 3.62 |
| 8 | 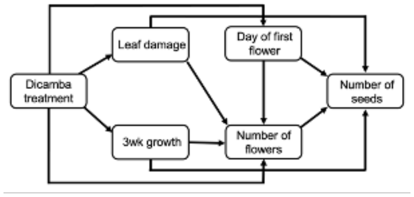 | 62.67 | 4.78 |
| 9 | 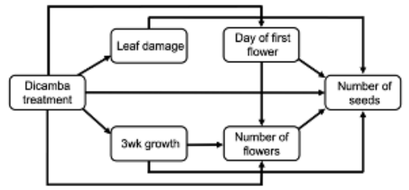 | 65.25 | 7.36 |
| 10 | 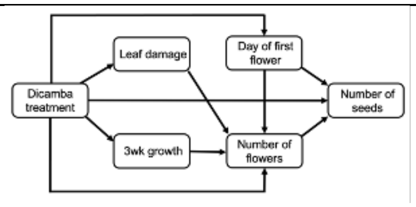 | 66.12 | 8.23 |
| 11 | 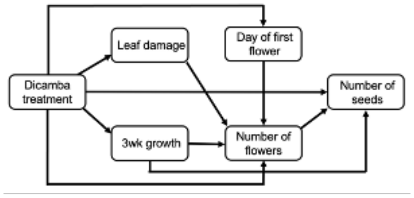 | 116.31 | 58.42 |
| 12 | 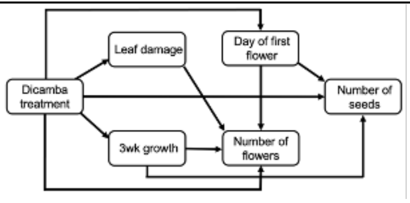 | 127.17 | 69.28 |
| 13 | 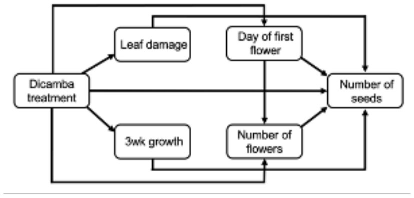 | 157.06 | 99.17 |
